# A chill brain-music interface for enhancing music chills with personalized playlists

**DOI:** 10.1101/2024.11.07.621657

**Authors:** Sotaro Kondoh, Takahide Etani, Yuna Sakakibara, Yasushi Naruse, Yasuhiko Imamura, Takuya Ibaraki, Shinya Fujii

## Abstract

Music chills are pleasurable experiences while listening to music, often accompanied by physical responses, such as goosebumps^1,2^. Enjoying music that induces chills is central to music appreciation, and engages the reward system in the brain^3–5^. However, the specific songs that trigger chills vary with individual preferences^6^, and the neural substrates associated with musical rewards differ among individuals^7–9^, making it challenging to establish a standard method for enhancing music chills. In this study, we developed the Chill Brain-Music Interface (C-BMI), a closed-loop neurofeedback system that uses in-ear electroencephalogram (EEG) for song selection. The C-BMI generates personalized playlists aimed at evoking chills by integrating individual song preferences and neural activity related to music reward processing. Twenty-four participants listened to both self-selected and other-selected songs, reporting higher pleasure levels and experiencing more chills in their self-selected songs. We constructed two LASSO regression models to support the C-BMI. Model 1 predicted pleasure based on the acoustic features of the self-selected songs. Model 2 classified the EEG responses when participants listened to self-selected versus other-selected songs. Model 1 was applied to over 7,000 candidate songs, predicting pleasure scores. We used these predicted scores and acoustic similarity to the self-selected songs to rank songs that were likely to induce pleasure. Using this ranking, four tailored playlists were generated. Two playlists were designed to augment pleasure by selecting top-ranked songs, one of which incorporated real-time pleasure estimates from Model 2 to continuously update Model 1 and refine song rankings. Additionally, two playlists aimed to diminish pleasure, with one updated using Model 2. We found that the pleasure-augmenting playlist with EEG-based updates elicited more chills and higher pleasure levels than pleasure-diminishing playlists. Our results indicate that C-BMI using in-ear EEG data can enhance music-induced chills.

---

Humans enjoy music universally across diverse cultures and societies^10,11^. Although the evolutionary functions of music remain controversial^12–14^, it is widely acknowledged that music evokes positive emotions accompanied by bodily sensations^6,15–19^. One such emotion is “music chills,” an intense pleasurable sensation accompanied by physiological responses such as “goosebumps” or a “shiver down the spine”^1,2^. Studies have shown that experiencing music chills correlates with sympathetic nervous responses, such as elevated skin conductance responses (SCR), heart rates, and increasing pupil diameters^20–23^, suggesting a strong association with emotional arousal.

Neuroimaging studies have demonstrated that experiencing music chills activates the brain’s reward system. For example, music chills increase blood flow to areas including the ventral striatum, anterior cingulate cortex, and insula, which are also activated by rewarding stimuli, such as food, sex, and drugs^3^. Further research indicates that the dorsal and ventral striatum (i.e., the caudate nucleus and nucleus accumbens) are involved in anticipating and experiencing music chills, respectively^4^. Moreover, when participants perceived the reward value of the music they listened to for the first time, the functional connectivity between the nucleus accumbens and auditory cortex strengthened^5^. These studies indicate that music chills involve both the reward and auditory perceptual systems^6^. Research measuring electroencephalogram (EEG) activity suggests that this complex interaction is represented by theta activity^24–26^. These pleasurable experiences result from positive reward prediction errors that arise when the perceived sound is better than expected^6,27–30^.

Researchers have attempted to enhance music chills by stimulating the reward system of the brain. For instance, excitatory intermittent theta-burst transcranial magnetic stimulation of the left dorsolateral prefrontal cortex elicits more pleasure than inhibitory continuous theta-bursts^31^. Taking a dopamine agonist as a pharmacological approach increased subjective music chills and pleasure more than a dopamine antagonist did, without a significant difference in SCR^32^. Meanwhile, an opioid agonist increased SCR without modulating subjective chills^33^, suggesting that dopamine regulates subjective responses, whereas opioids regulate physiological responses to musical rewards. However, individual differences in musical preferences and sensitivity to musical rewards present a challenge for standardized methods to enhance chills. Functional connectivity between the auditory cortex and reward-related areas suggests various musical preferences, as individuals have different auditory templates based on their music exposure and cultural or social influence^5,6^. The Barcelona Music Reward Questionnaire (BMRQ) revealed individual variability in musical reward sensitivity and its neural substrates^7–9^. Thus, developing methods to improve music chills requires more information about individuals’ musical preferences and their neural responses to music pleasure.

One promising approach is closed-loop neuromodulation, which adjusts the presented stimuli based on neural activity to guide it toward a specific state. For example, implanting electrodes in the deep cortex of a depressed patient and providing personalized electrical stimulation improves the symptoms^34^. In the neuroscience of music, visualizing listeners’ amygdala activity in real time and presenting it to a professional pianist, who then maximizes it through live performance, increases neural activity in regions such as the amygdala, hippocampus, and ventral striatum, resulting in higher emotional arousal and valence than recorded music^35^. As a closed-loop neuromodulation system adjusts and optimizes stimuli for individuals, it can overcome individual differences. In the present study, we examined the effect of the “Chill Brain-Music Interface (C-BMI),” a closed-loop song selection system based on neural activity optimized for each individual to induce music chills. We hypothesized that a song playlist designed to increase pleasure using EEG would enhance chills more than other playlists would.

## C-BMI overview

The C-BMI consists of three steps (Fig.1):

**Fig.1.**
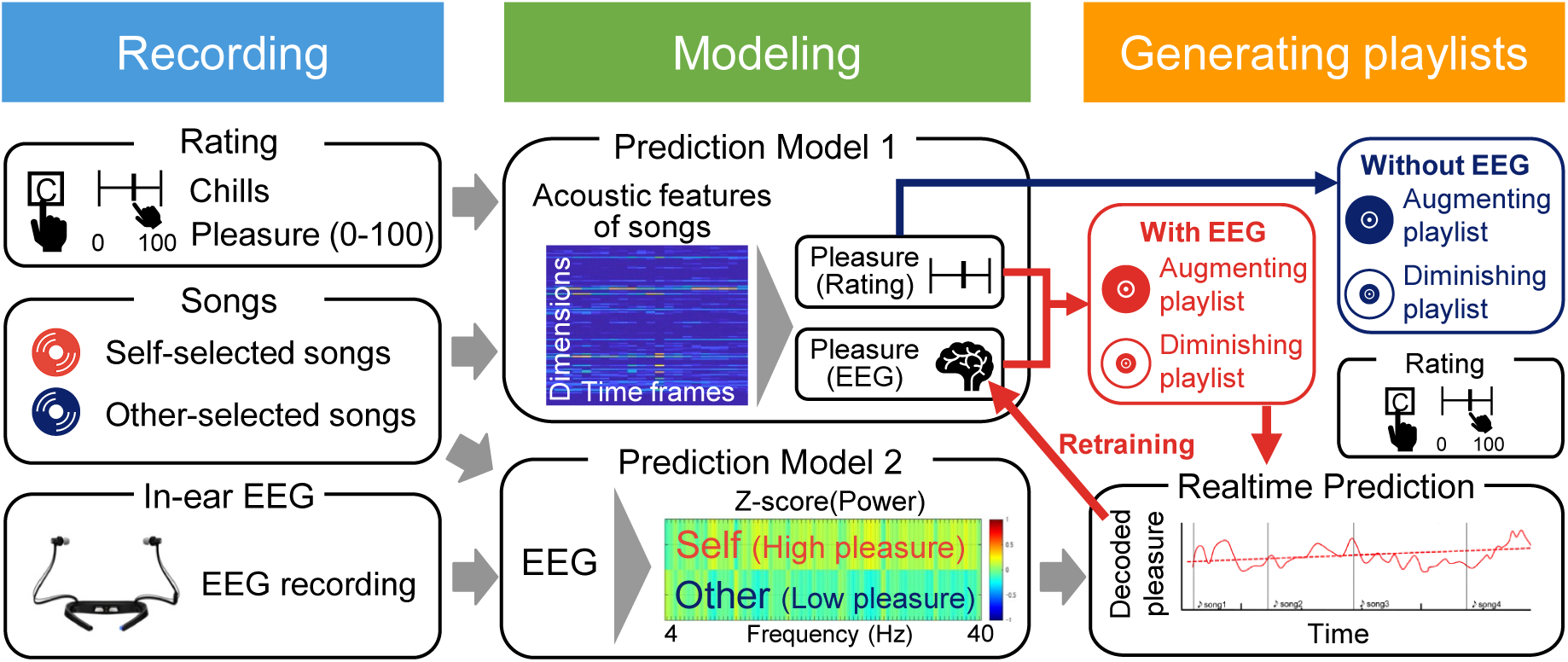
Overview of the “Chill Brain-Music Interface (C-BMI).” First, participants listened to three self-selected songs and three other-selected songs. They pressed a key when experiencing music chills and rated the pleasure level of each song using a visual analog scale (VAS) after listening to it. Their in-ear electroencephalogram (EEG) was recorded while listening to songs. Second, we created two pleasure prediction models. Model 1 predicted pleasure based on the acoustic features of the songs. Model 2 classified the EEG state when participants listened to self-selected (high pleasure) or other-selected (low pleasure) songs, enabling pleasure prediction from in-ear EEG. Third, tailor-made song playlists were generated. We used Model 1 to predict the pleasure level from candidate songs, added acoustic similarity to self-selected songs, and sorted them in descending order to make a song ranking that was likely to induce pleasure. The pleasure-augmenting playlist updated using EEG selected songs from the top of the ranking. Model 2 decoded real-time pleasure levels from the in-ear EEG while participants listened to each song on the playlist. The decoded pleasure levels and acoustic features of the song were used to retrain Model 1 and update the song rankings. The pleasure-diminishing playlist with EEG updates used the same procedure, but selected songs from the bottom of the ranking. Augmenting and diminishing playlists without EEG selected songs from the top or bottom of the ranking without using Model 2 and did not retrain Model 1. Participants pressed a key when experiencing chills, and rated the emotion and well-being items for each playlist using a VAS after listening to each playlist.

**Recording:** We obtained chill-inducing songs from each participant before the experiment and measured their in-ear EEG data while they listened to their chosen songs. Twenty-four participants listened to three songs that had previously induced chills, and three additional songs selected by another participant. The duration of each song was 90 seconds. Participants pressed a key whenever they experienced chills while listening to each song and rated their pleasure levels using a visual analog scale (VAS: 0-100) at the end of listening. In-ear EEG data were recorded throughout each song presentation.

**Modeling:** We confirmed whether participants experienced more chills and higher pleasure with self-selected songs. We analyzed the acoustic features of self-selected songs using VGGish^36,37^, a pretrained neural network, and developed a linear LASSO model to predict subjective pleasure levels from acoustic features (Model 1). Additionally, we created a generalized LASSO model to classify EEG states associated with high pleasure (while listening to self-selected songs) and low pleasure (while listening to other-selected songs) (Model 2).

**Generating playlists:** We generated four tailor-made music playlists for each participant to augment or diminish pleasure with or without the EEG. The playlists were named (1) ‘augmenting playlist with EEG (AugEEG),’ (2) ‘augmenting playlist without EEG (AugNoEEG),’ (3) ‘diminishing playlist with EEG (DimEEG),’ and (4) ‘diminishing playlist without EEG (DimNoEEG).’ We computed the acoustic features of 7,225 candidate songs using VGGish, and sorted them based on the pleasure scores predicted by Model 1 and their acoustic similarity to self-selected songs to create a ranking of songs that are likely to induce pleasure. For Playlists (1) and (2), songs were randomly chosen from the top ranking, whereas for Playlists (3) and (4), songs were chosen from the bottom ranking. The first song in each playlist was randomly selected from the middle range of the ranking (positions 3,577–3,649) to serve as the baseline for analysis. Each playlist comprised seven songs, and the duration of each song was 90 seconds. The in-ear EEG was recorded while the participants listened to songs. For the playlists with EEG updates (1 and 3), the decoded pleasure from the EEG using Model 2 and the acoustic features of the song using VGGish were used to retrain Model 1 and update the song ranking. The playlists without EEG updates (2 and 4) did not retrain Model 1 or update the rankings. The participants listened to all the playlists. They pressed the key when experiencing chills while listening to songs and rated items related to emotions and well-being, such as the subjective pleasure level, valence, arousal, and preference (Table 1) of the entire playlist using a VAS (0-100) after listening to all songs in each playlist.

**Table 1.** Subjective rating items.

| Label | Item |
| --- | --- |
| Pleasure | I felt pleasure in the whole playlist. |
| Excitement | I felt excited throughout the playlist. |
| Absorption | I became absorbed while listening to this playlist. |
| Arousal | I am currently aroused. |
| Valence | I am currently in a comfortable state. |
| Preference | I prefer this playlist. |
| Strangeness | I feel strange about this playlist. |
| Life purpose | Listening to this playlist helped me feel more connected to the purpose and meaning of life. |
| Self-acceptance | Listening to this playlist helped me accept myself as I am. |
| Improving relations | Listening to this playlist helped me improve relationships with others. |
| Reducing stress | Listening to this playlist helped me feel less stressed and more relaxed. |
| Resilience | Listening to this playlist helped me think about dealing more effectively with difficult situations. |
| Emotion regulation | Listening to this playlist helped me better understand and accept my emotions and feelings. |
| Seeking help | Listening to this playlist helped me think about finding support and solutions to the problems and difficulties I am facing. |

## Chill counts

Chill counts for the four playlists (Fig.2A) were as follows: AugEEG had the highest count (10.50 ± 9.52; Mean ± SD), followed by AugNoEEG (9.50 ± 12.15), DimNoEEG (6.25 ± 7.64), and DimEEG (4.90 ± 6.37). The normality of the chill counts for all playlists was not confirmed (Extended Data Table 1). The Friedman test revealed a significant difference in the number of chills across the playlists (*χ^2^*= 12.81, *p* = 0.005, *η^2^* = 0.21). Paired Wilcoxon signed-rank tests revealed that the chills of AugEEG were significantly more than DimEEG (*V* = 179.5, *p_Holm_* < 0.001). However, there were no significant differences between AugEEG and DimNoEEG (*V* = 156.5, *p_Holm_* = 0.055) or AugEEG and AugNoEEG (*V* = 138.5, *p_Holm_* = 0.242). Additionally, AugNoEEG did not show a significant difference compared to DimEEG (*V* = 128.5, *p*_Holm_ = 0.055) or DimNoEEG (*V* = 123.0, *p_Holm_* = 0.542). DimNoEEG was not significantly different from DimEEG (*V* = 68.5, *p_Holm_* = 0.656). These results suggest that AugEEG induces more chills than DimEEG.

**Fig.2.**
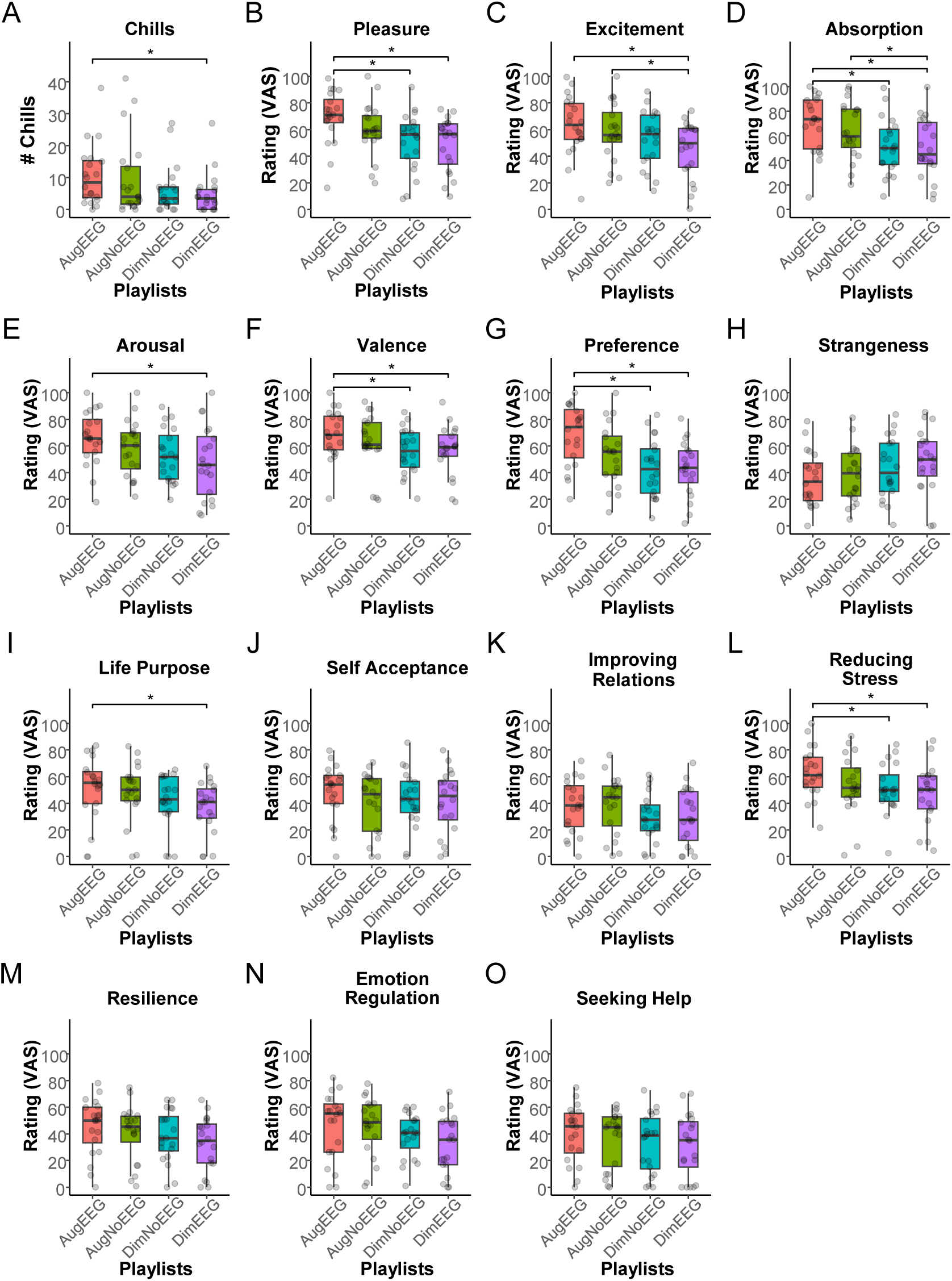
Boxplots showing the number of chills and subjective ratings (*n* = 20). Each dot represents the value for each participant. **A.** Number of chills (# chills) for each playlist. The augmenting playlist with EEG (AugEEG) elicited a significantly larger number of chills than the diminishing playlist with EEG (DimEEG). **B.** Subjective pleasure ratings. AugEEG received significantly higher pleasure ratings than the diminishing playlist without EEG (DimNoEEG) and DimEEG. Note that AugNoEEG means the augmenting playlist without EEG. **C-O.** Subjective ratings of excitement, absorption, arousal, valence, preference, strangeness, life purpose, self-acceptance, improving relations, reducing stress, resilience, emotion regulation, and seeking help. Subjective ratings were assessed using the VAS. The *p*-values were adjusted using Holm’s method. Asterisks (*) indicate significant differences (*p*_Holm_ < 0.05).

## Subjective pleasure rating

The subjective pleasure ratings (VAS) for the four playlists (Fig.2B) were as follows: AugEEG had the highest rating (69.47 ± 20.04; Mean ± SD), followed by AugNoEEG (60.05 ± 19.60), DimNoEEG (51.06 ± 21.90), and DimEEG (49.17 ± 20.05). We confirmed the normality of ratings for all playlists (Extended Data Table 1). One-way repeated-measures ANOVA showed that pleasure levels differed significantly across playlists (*F*(3, 57) = 6.31, *p* < 0.001, *η_p_^2^* = 0.25, Greenhouse-Geisser (GG) *ε* = 0.82). The paired *t*-tests revealed that the pleasure of AugEEG was significantly higher than that of DimEEG (*t*(19) = 4.17, *p*_Holm_ = 0.003, 95% confidence interval (CI) [10.12, 30.48]) and DimNoEEG (*t*(19) = 3.89, *p*_Holm_ = 0.005, 95%CI [8.50, 28.33]). However, there were no significant differences between AugEEG and AugNoEEG (*t*(19) = 1.51, *p*_Holm_ = 0.443, 95%CI [−3.64, 22.47]). Similarly, AugNoEEG did not show a significant difference compared to DimEEG (*t*(19) = 2.00, *p*_Holm_ = 0.239, 95%CI [−0.50, 22.27]) or DimNoEEG (*t*(19) = 1.47, *p*_Holm_ = 0.443, 95%CI [−3.84, 21.83]). Finally, DimNoEEG was not significantly different from DimEEG (*t*(19) = 0.52, *p*_Holm_ = 0.609, 95%CI [−5.70, 9.48]). These results indicate that AugEEG evokes more pleasure than DimEEG or DimNoEEG.

## Subjective ratings of items related to emotions and well-being

One-way repeated-measures ANOVAs or Friedman tests revealed significant differences among playlists for the following items (Fig.2C-O): excitement (*F*(3, 57) = 4.03, *p* = 0.011, *η_p_^2^* = 0.17, GG *ε* = 0.78), absorption (*F*(3, 57) = 6.85, *p* < 0.001, *η_p_^2^* = 0.27, GG *ε* = 0.86), arousal (*F*(3, 57) = 4.90, *p* = 0.010 (GG corrected), *η_p_^2^* = 0.21, GG *ε* = 0.74), valence (*χ^2^* = 14.02, *p* = 0.003, *η^2^* = 0.23), preference (*F*(3, 57) = 9.78, *p* < 0.001, *η_p_^2^* = 0.34, GG *ε* = 0.88), life purpose (*χ^2^* = 8.85, *p* = 0.031, *η^2^*= 0.15), and reducing stress (*F*(3, 57) = 3.36, *p* = 0.025, *η_p_^2^* = 0.15, GG *ε* = 0.80). Subsequent multiple comparisons (paired *t*-tests or paired Wilcoxon signed-rank tests) revealed that the values of excitement (*t*(19) = 3.01, *p_Holm_* = 0.036), absorption (*t*(19) = 3.57, *p_Holm_* = 0.010), arousal (*t*(19) = 3.04, *p_Holm_* = 0.041), valence (*V* = 179, *p_Holm_* = 0.021), preference (*t*(19) = 5.40, *p_Holm_* < 0.001), life purpose (*V* = 147, *p_Holm_* = 0.034), and reducing stress (*t*(19) = 3.50, *p_Holm_* = 0.012) were higher in AugEEG than in DimEEG. In addition, the ratings of AugEEG were higher than those of DimNoEEG for the following items: absorption (*t*(19) = 3.75, *p_Holm_* = 0.008), valence (*V* = 171, *p_Holm_* = 0.007), preference (*t*(19) = 4.96, *p_Holm_* < 0.001), reducing stress (*t*(19) = 3.64, *p_Holm_* = 0.011). Moreover, the excitement (*t*(19) = 3.15, *p_Holm_* = 0.032) and absorption (*t*(19) = 2.88, *p_Holm_* = 0.038) of AugNoEEG were higher than those of DimEEG. However, no significant differences were found among the playlists for the following items: strangeness (*F*(3, 57) = 1.89, *p* = 0.141, *η_p_^2^* = 0.09, GG *ε* = 0.83), self-acceptance (*F*(3, 57) = 0.92, *p* = 0.404 (GG corrected), *η_p_^2^* = 0.05, GG *ε* = 0.64), improving relations (*F*(3, 57) = 2.35, *p* = 0.082, *η_p_^2^* = 0.11, GG *ε* = 0.76), resilience (*χ^2^*= 5.75, *p* = 0.124, *η^2^* = 0.10), emotion regulation (*F*(3, 57) = 2.63, *p* = 0.059, *η_p_^2^* = 0.12, GG *ε* = 0.79), and seeking help (*χ^2^* = 3.35, *p* = 0.341, *η^2^* = 0.06). See Extended Data Table 1-8 for the mean and standard deviation of the subjective evaluation items and the differences (including 95% CI) between playlists.

## Decoded pleasure from in-ear EEG

To examine whether the pleasure levels decoded from the in-ear EEG differed across playlists, we constructed a Bayesian generalized linear mixed-effects model (GLMM) by putting the baseline-corrected decoded pleasure from the in-ear EEG into the response variable, the number of songs and the playlists into fixed effects, and participant ID into the random effect (Extended Data Table 9). The baseline was defined as the first 90 seconds of each playlist, corresponding to the first song. The slope of the song (the number of songs) was negative (estimate = −0.009, 95% CI [−0.013, −0.005]), suggesting that as songs progressed in the playlist, the pleasure level decoded from the EEG decreased. After removing the decreasing trend, AugEEG showed the highest decoded pleasure, with an increasing trend from the baseline, followed by AugNoEEG, DimNoEEG, and DimEEG (Fig.3A). Pairwise comparisons between the levels of playlists (Fig.3B, Extended Data Table 10) revealed a difference between AugEEG and DimEEG (estimate = 0.034, 95% Highest Posterior Density (HPD) intervals [0.011, 0.056]), with AugEEG having higher decoded pleasure values, which is consistent with the results for chills and the VAS of pleasure. Another difference was also found between AugEEG and AugNoEEG (estimate = 0.038, 95% HPD Intervals [0.015, 0.060]), indicating that AugEEG having higher decoded pleasure values than AugNoEEG. The 95% HPD intervals for the other contrasts did not include zero.

**Fig. 3.**
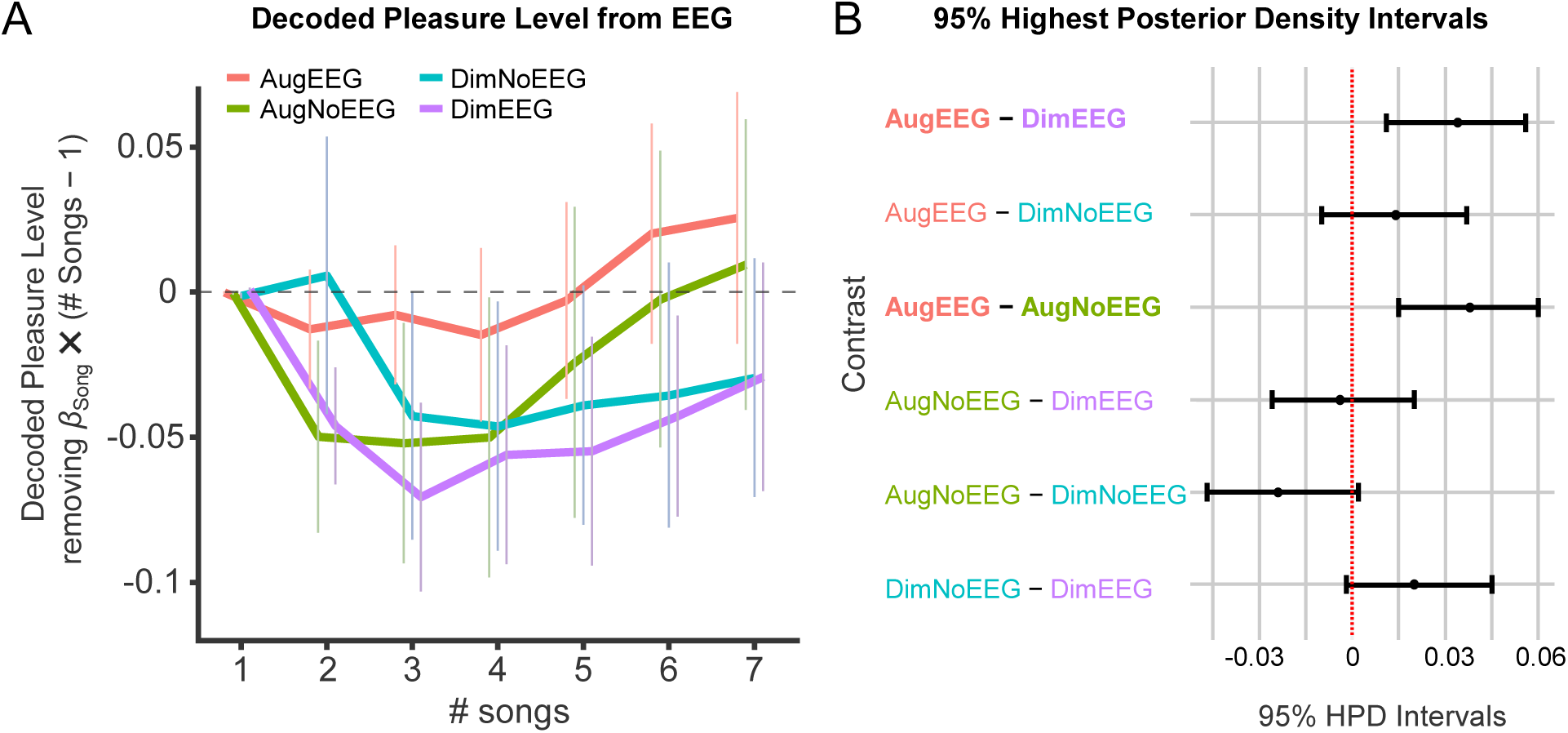
Decoded pleasure levels from in-ear EEG and pairwise comparisons (*n* = 17). **A.** Mean decoded pleasure level from in-ear EEG. The Bayesian generalized linear mixed-effects model (GLMM) was applied as follows: *Decoded pleasure ∼ Song (number of songs) + Playlist + (1 | Participants ID)*. The effects of the songs (*β*_Song_ × (*Song* − 1)) were removed from the plot. The pleasure level of the AugEEG increased from the baseline (the first song) as the playlist progressed. Error bars indicate the standard error. **B.** 95% highest posterior density intervals of pairwise comparisons. The difference between AugEEG and DimEEG did not include zero, with AugEEG having higher decoded pleasure values than did DimEEG. The difference between AugEEG and AugNoEEG also did not contain zero, with AugEEG having a higher decoded pleasure than AugNoEEG.

## Effects of music reward sensitivity and musical sophistication

To examine whether individual differences in music reward sensitivity and musical sophistication influence chills and pleasure derived from the playlists, we collected participants’ scores on the Japanese version of the BMRQ (J-BMRQ)^7,38^ and the Goldsmiths Musical Sophistication Index (Gold-MSI)^39,40^. We performed multiple regression analyses to explore the factors within the J-BMRQ and Gold-MSI that affect the overall pleasure derived from the playlists. We calculated differences in chills (Chilldiff) and subjective pleasure (Pldiff) between the two EEG playlists (AugEEG minus DimEEG). We then calculated the z-scores of the Chilldiff and Pldiff and summed them to obtain the total pleasure score for the EEG playlists (total pleasure EEG). We created a model with a response variable as the total pleasure EEG and all factors of the J-BMRQ and Gold-MSI as explanatory variables. A stepwise regression approach was used to determine the final model (*F*(3,13) = 5.73, *p* = 0.010, Adjusted *R^2^*= 0.47; Extended Data Table 11). The VIFs of all selected variables were less than 2. The normality of the residuals was confirmed (Shapiro-Wilk test: *W* = 0.97, *p* = 0.761). The results suggest that listeners with more musical training and emotion factor scores experience more pleasure in AugEEG than in DimEEG, although higher singing ability scores connect less pleasure in AugEEG.

We also calculated the total pleasure score by subtracting the pleasure score of DimNoEEG from that of AugNoEEG (total pleasure NoEEG). A multiple regression model was constructed with the total pleasure NoEEG score as the response variable, a similar stepwise regression approach was utilized, and the final model was determined (*F*(3,13) = 5.18, *p* = 0.014, Adjusted *R^2^* = 0.44; Extended Data Table 12). The VIFs of all selected variables were less than 2. The normality of the residuals was confirmed (Shapiro-Wilk test: *W* = 0.98, *p* = 0.983). This result indicates that participants with more mood regulation factors experienced less pleasure in AugNoEEG than in DimNoEEG.

## Discussion

This study aimed to determine whether selecting music based on pleasure decoded from in-ear EEG would increase chills and pleasure. AugEEG resulted in the highest number of chills among the playlists, significantly surpassing DimEEG (Fig.2A). Similarly, subjective pleasure ratings were significantly higher for AugEEG than for DimEEG and DimNoEEG (Fig.2B). These findings suggest that personalized song playlists created using C-BMI, a closed-loop neurofeedback system with in-ear EEG, can elicit more chills and greater pleasure. Previous studies have shown that electrical stimulation based on neural activity related to depression symptoms can improve these symptoms when data are collected using electrodes implanted in the deep cortex^34^. Additionally, live piano performance tailored to the participants’ amygdala activity can enhance its neural activity^35^. Our results are consistent with these findings and demonstrate the effectiveness of closed-loop neurofeedback, demonstrating that this approach can enhance music-induced chills. The method we employed showed the potential to induce chills in a simpler and less burdensome way for listeners compared to previous studies^31,32^ that used TMS or dopamine agonists to enhance the pleasure of music.

In this study, we collected ratings related to emotions and well-being for each playlist (Fig.2C-O). The ratings of AugEEG were higher than those of DimEEG, in terms of excitement, absorption, arousal, valence, and preference. Conversely, AugNoEEG, which did not use EEG, did not produce significantly higher values than the other playlists except for excitement and absorption. These results suggest that updating playlists using an individual’s EEG can effectively lead to excitement, absorption, and arousal and is preferred by individuals. Current popular music distribution services, such as Spotify, recommend songs based on users’ listening history^41^, similar to the creation of our AugNoEEG playlists. However, our results indicate that it is possible to create preferred playlists using neurofeedback. Interestingly, AugEEG had a higher life purpose score than DimEEG and a higher stress reduction score than both DimEEG and DimNoEEG (Fig.2I,L). These results indicate that neurofeedback can enhance well-being by helping listeners reaffirm their meaning and purpose in life and reduce their stress. Previous studies have shown that music can improve well-being, especially during periods of isolation due to COVID-19^42,43^. By applying our findings, we propose new methods for listening to music and creating playlists that can further improve well-being using an individual’s EEG.

Using Model 2 to estimate pleasure levels from EEG, we found that AugEEG values were higher than DimEEG values (Fig.3A,B). This aligns with the increased number of music chills and higher subjective pleasure ratings observed for AugEEG compared to DimEEG (Fig.2A,B), supporting the validity of decoding pleasure from EEG data. Additionally, the decoded pleasure value for AugEEG was greater than that for AugNoEEG (Fig.3A,B), suggesting that incorporating an individual’s EEG may enhance pleasure more effectively than playlists based solely on preferences. However, because we employed a two-channel in-ear EEG, the source localization was not feasible. As a result, we could not determine whether the EEG signals originated from the nucleus accumbens or other reward-related brain regions. Previous studies have demonstrated that it is possible to estimate nucleus accumbens activity during music listening by combining EEG with functional magnetic resonance imaging^44^. Therefore, future studies should address which brain areas in-ear EEG signals originate from during music listening.

In this study, we used the J-BMRQ^7,38^ and the Japanese version of the Gold-MSI^39,40^ to assess the effects of participants’ music reward sensitivity and musical sophistication on music chills and pleasure elicited by playlists. The chills and pleasure ratings for AugEEG and DimEEG were converted into z-scores and summed, and a stepwise regression was used to identify which of the 10 factors of music reward sensitivity and music sophistication predicted the total pleasure. The results showed that the slopes of the music training and emotion factors were significantly positive, whereas the slope of the singing ability factor was significantly negative (Extended Data Table 11). This indicates that greater music training or higher responsiveness to music emotions correlates with increased pleasure from AugEEG compared to DimEEG. The positive slope of the music-training factor is consistent with recent studies that reported that music training modulates emotional responses to music^45,46^. The emotion factor included questions like “I sometimes choose music that can trigger shivers down my spine,” which likely explains its amplification of the pleasure from AugEEG. Conversely, higher subjective singing ability was associated with greater pleasure from the DimEEG. Individuals with better singing abilities may listen to songs more analytically, making it more difficult to evoke emotions. In a similar stepwise regression model using the difference in total pleasure between AugNoEEG and DimNoEEG as the response variable, mood regulation, sensorimotor, and perception ability were selected as explanatory variables, with only mood regulation being significant and having a negative slope (Extended Data Table 12). This means that higher emotion regulation is associated with greater pleasure from the DimNoEEG. These models suggest that playlists with EEG updates reflect more individual differences in musical experiences than playlists without EEG updates, supporting the idea that playlists with EEG updates are more personalized in eliciting pleasure.

This study has several limitations. First, we did not measure autonomic nervous system activity during playlist listening; therefore, we could not confirm whether physical sensations such as goosebumps actually occurred. Future studies should address this issue; however, given that previous research has shown that arousal-related autonomic responses increase during chills^23^ and that arousal rating was also high for AugEEG in this study, it is likely that goosebumps occurred. Second, the GLMM fitted to the decoded pleasure levels indicated a significantly negative slope for the number of songs on the playlist. This suggests that, as the number of songs increased, the overall decoded pleasure level decreased. Considering that it took 40 minutes to listen to all the playlists, with each song lasting 90 seconds, it is possible that fatigue from the long measurement and changes in ear conditions affected the decoded pleasure value. Future studies should clarify this decreasing trend and measure pleasure after eliminating these effects. Finally, in AugEEG and DimEEG, Model 1 was retrained using pleasure estimation from the EEG after each song. In future studies, it may be possible to update the model using other indices, such as subjective ratings or autonomic nervous system activity. Continuous subjective ratings and autonomic nervous system activity, such as the heart rate, are candidates for these indices. Future research should explore and compare alternative methods for updating models using EEG data.

## Conclusion

This study demonstrated that music chills and pleasure can be amplified by estimating pleasure from an individual’s EEG and creating a tailor-made music playlist based on the data. Personalized playlists can provide insights into the mechanisms that induce and augment musical rewards, and they have the potential to improve listeners’ well-being by helping people reaffirm the meaning and purpose of their lives through music and reducing stress. Although future research should measure autonomic nervous system activity during playlist listening to verify its effects on the body and improve the accuracy of pleasure decoding, we believe that C-BMI paves the way for personalizing music, enhancing pleasure from music, and offering new technology for a more enjoyable music experience.

## Methods

### Participants

We recruited twenty-four participants (10 males, 14 females; 23.8 ± 6.5 years old) and obtained written informed consent from all participants. All experimental procedures were approved by the Research Ethics Committee of Keio University Shonan Fujisawa Campus, Japan (#524). The experiment was conducted from February 14, 2024, to May 3, 2024. The sample size of 24 participants was determined beforehand using G*Power 3 software for one-way repeated-measures analysis of variance (ANOVA), assuming a medium-sized effect, *α* = 0.05, and *power* = 0.8^47^.

### Recording

#### Self-selected and other-selected songs

Participants were informed that music chills are “intense and pleasurable sensations accompanied by goosebumps while listening to music” and were asked whether they had experienced the sensations. After confirming these experiences, each participant selected three songs (including instrumentals) during which they personally experienced music chills multiple times. The participants provided songs as YouTube links or mp3 files to the first author (the experimenter) before the experiment day. They also specified particular moments within the songs when they were most likely to experience chills. See Extended Data Table 13 for song information.

#### In-ear EEG with the number of chills and pleasure ratings

Participants listened to three self-selected songs and three songs selected by another participant in random order. The assignment of songs selected by another participant was random, with no duplicates between participants. The playback time for each song was 90 seconds, which included the parts where participants frequently experienced chills. If no specific time was reported, the first 90 seconds of the song was played. The participants adjusted their volume to a comfortable level before the playback.

While listening to the songs, we measured the participants’ in-ear EEG using a VIE CHILL device (VIE Inc., Japan). EEG data were recorded from the left and right ear canals with a reference electrode placed on the back of the neck. We attached electrically conductive ear tips to two electrodes inserted into the ear canals. These electrodes also function as earphones, allowing for simultaneous song playback and EEG collection. The sampling rate was set to 600 Hz. Participants pressed the C key on the keyboard when they experienced chills. To avoid noise from other physical movements, such as electromyography, the participants were instructed to remain relaxed and avoid moving, except when pressing the key. At the end of each song, the participants rated their level of pleasure on a visual analog scale (VAS) from 0 to 100. After the evaluation, the next song was played and the EEG recording was restarted.

### Modeling

#### Model 1: A pleasure predicting model from acoustic features

We developed a linear LASSO model to predict pleasure levels from the acoustic features of the songs that participants listened to. The acoustic features were calculated using VGGish^36,37^, which is a pretrained neural network. For each participant, VGGish transformed six 90-second song excerpts (three self-selected and three other-selected) into spectrograms and calculated 372 time frames (determined based on 90-second duration) and 128 feature dimensions. The values of the acoustic features were averaged for each time frame corresponding to 10 seconds, resulting in a matrix of 6 songs × 9 means × 128 dimensions.

We calculated the mean and standard deviation of the features across the six excerpts for each dimension, and computed the z-scores of the acoustic features. Additionally, we adjusted the z-scores by selecting dimensions where the mean was not a missing value (NaN) and the standard deviation was greater than 0.01 to improve the accuracy of the model. Finally, we constructed a linear LASSO model with the z-scores of the pleasure ratings as the response variable and the adjusted z-scores of the acoustic features as the explanatory variables as

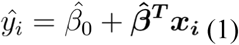

where 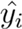 is the predicted subjective pleasure rating for song *i*. 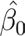 is the intercept, and 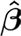 is a vector of coefficients for each acoustic feature dimension. *x_i_* is a vector of selected acoustic feature dimensions calculated by VGGish for song *i*. Note that 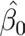 and 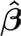 are the variables of *β*_0_ and *β* that minimize the following loss function:

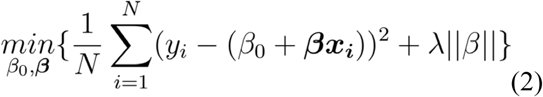

where *λ* is the regularization parameter. The optimal *λ* was selected using 5-fold cross-validation. *N* represents 6, the number of the songs.

#### Model 2: A pleasure predicting model from in-ear EEG

To develop a model classifying EEG signals into states of high or low pleasure, we tested whether participants experienced more chills and a higher level of pleasure with the songs they chose themselves than with songs chosen by another participant (Extended Data Fig.1A,B). The criterion for chills was that they experienced three or more chills in self-selected songs than in other-selected songs, based on previous research^48^. Furthermore, the criterion for pleasure was that the average level of the three self-selected songs should be higher than that of songs chosen by others. Two participants did not meet these inclusion criteria.

For preprocessing the EEG data, the raw data were filtered using a fourth-order Butterworth filter with a frequency range of 3–40 Hz. We performed an independent component analysis (ICA) using filtered signals from the left and right electrodes and extracted two components. We calculated the RMS of the two components, labeled the component with the larger RMS as noise, and removed it. We set a time window of four seconds with a 50% overlap. We applied a time window to the EEG data and extracted the epochs. For each epoch, we calculated the power spectral density from 4 to 40 Hz at 0.5 Hz intervals. We also calculated the sum of the powers across all frequency bands and obtained the RMS of the filtered data for the two channels (left and right ears). In addition, we obtained the maximum value of the numerical gradient of the signal and the skewness in the same manner. Each noise metric was standardized and averaged for the right and left electrodes and each time window, and a noise flag was set for data that exceeded the predetermined threshold of 2.5. We used only epochs without noise flags.

The EEG data were standardized for all epochs, frequency windows, and electrodes (left, right, and their differences). Principal component analysis (PCA) was performed on the standardized EEG data. The maximum number of principal components was 150. If the actual number of principal components was greater than 150, then the number was adjusted to 150. If the actual number of principal components was less than 150, all the components were used. The mean and standard deviation of the principal component scores were calculated and standardized. Each epoch was categorized based on whether it was recorded while listening to self-selected (true) or other-selected songs (false). If the number of true and false data points was uneven, the data were randomly extracted from the epoch with more data to match a smaller number. Ninety percent of the data were used for training and ten percent for testing.

We constructed a logistic LASSO regression model to decode pleasure based on EEG frequency components. The model’s prediction for the odds of pleasure, denoted as 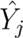, is expressed as

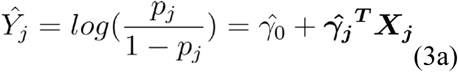

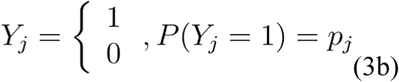

where *p_j_* is the probability that the EEG data are classified into a state when listening to self-selected songs, *j* is the number of EEG components, 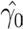 is the intercept, and 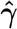 is a vector of coefficients corresponding to each EEG component. *X_j_* represents a vector of the EEG components. Note that *Y_j_* is the binary outcome variable. 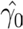 and 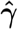 are the variables of *γ*_0_ and *γ* that minimize the following loss function:

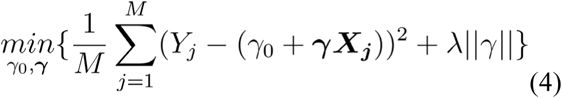

where *λ* is the regularization parameter. The optimal *λ* was selected using 20-fold cross-validation. *M* represents the maximum value of the EEG components.

We then calculated the confusion matrix, including true positives, false positives, false negatives, and true negatives, and the accuracy rate of the training data, as well as the receiver operating characteristic (ROC) curve and the area under the curve (AUC) (Extended Data Fig.1C). The model was trained using the training data and evaluated using the test data by performing the same prediction and evaluation on the test data. The ROC curve and the area under the curve (AUC) for the test data were also calculated. The mean accuracy rates for the trained and test data were 83.6% and 73.6%, respectively (Extended Data Fig.1D,E), and the mean AUC for the trained and test data were 0.90 and 0.80, respectively (Extended Data Fig.1F,G). As analyzed earlier, participants experienced higher subjective pleasure with the songs they selected and lower pleasure with songs selected by others. Hence, this generalized logistic LASSO model that classifies true and false epochs can be used to estimate pleasure from EEG.

### Generating Playlists

#### Songs

The candidate songs for the playlists consisted of 7,225 tracks from the GfK Japan weekly music chart Top 1-1,000, spanning from April 2018 to February 2022. The feature values of these candidate songs were calculated using VGGish. These values were then normalized, with values exceeding two standard deviations being clipped. A matrix was created to predict the subjective pleasure level using Model 1. The scores were calculated by adding the z-scores of the correlation coefficients of the feature values of each candidate song to the participants’ self-selected songs.

Candidate songs were then ranked in descending order based on these scores. In-ear EEG measurement and decoding

EEG data were measured during playlist listening at a sampling rate of 600 Hz and a frequency range of 4-40 Hz. A Butterworth filter was applied to the signal in the 3-40 Hz band. The filtered EEG data were used to calculate the power spectrum, which was normalized and input into Model 2 to predict pleasure. For noise determination, metrics such as root mean square (RMS), maximum gradient, and kurtosis were calculated, and a noise flag was set. The criteria for removing noise were the same as those used in the previous modeling section. The moving average of the decoded pleasure state was calculated every 5 second, resulting in one data point per second. If the length of the decoded pleasure data was less than five, the average of the data points that were not flagged as noise was calculated. If it was five or more, the average of the latest five data points that were not flagged as noise was calculated.

#### Procedure

Each participant listened to four playlists consisting of seven songs. Each song lasted 90 seconds. The first song, which was used as the baseline, was randomly selected from positions 3,577 to 3,649. At the end of each song, 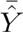, the average predicted pleasure for 90 seconds calculated by Model 2, excluding noise sections, was converted to an approximate value of a VAS, 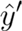, as follows:

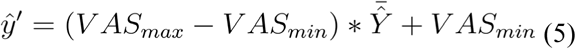

where *VAS_mzx_* and *VAS_min_* are the maximum and minimum VAS ratings in the first recording section, respectively. 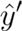 was standardized using the mean and standard deviation of the VAS ratings. The acoustic features of the music were precalculated using VGGish and converted to z-scores. We then added the standardized decoded pleasure level from the EEG and acoustic features of the played songs to Model 1 and retrained the model. This procedure was repeated until the playlist was complete.

The four playlists consisted of two pleasure-augmenting and two pleasure-diminishing playlists. Of the two augmenting playlists, AugEEG used the updated Model 1 to calculate the pleasure prediction score for the candidate songs. This score was then combined with the z-score of the correlation coefficient of the feature values with the participants’ self-selected songs, and the candidate songs were resorted in descending order based on this score to update the ranking. This playlist consisted of the top 72 songs (top 1% of all songs). The other augmenting playlist, AugNoEEG, was not updated using the retrained Model 1, but remained fixed using the initial ranking and was not updated using EEG. This playlist consisted of the top 432 songs (top 6% of all songs) to prevent any overlap between the two playlists. AugEEG repeated the selection of the top 72 songs six times; therefore, including duplicates, 432 songs were selected. To match this number, AugNoEEG had a selection range of 432 songs. The two diminishing playlists also included an EEG-updated playlist, DimEEG, and a non-updated playlist, DimNoEEG. DimEEG consisted of the bottom 72 songs (bottom 1% of all songs), whereas DimNoEEG consisted of the bottom 432 songs (bottom 6% of all songs).

Participants pressed the C key when they experienced chills while listening to each song. At the end of each playlist, they reflected on the entire playlist and rated it using 14 items (Table 1) using a VAS (0-100). After listening to all playlists, participants answered the Japanese version of the Barcelona Music Reward Questionnaire (J-BMRQ)^7,38^ and the Japanese version of the Goldsmiths Musical Sophistication Index (Gold-MSI)^39,40^.

### Statistics

All statistical analyses were performed using the R software version 4.1.1^49^. In the analysis of chill counts and VAS ratings, we excluded four participants and analyzed the data from 20 participants. Two participants did not experience three or more chills^48^ or higher pleasure from self-selected songs than from other-selected songs in the previous modeling section. Additionally, the pleasure level decoded from the EEG of one participant was consistently 0.9 or higher across the three playlists, indicating failure in decoding and feedback. The fourth participant was excluded because more than 30% of their EEG data during DimEEG was judged to be noise. We calculated the number of music chills and the VAS values for pleasure for each playlist. We hypothesized that AugEEG would induce more chills and higher pleasure than the other playlists (AugNoEEG, DimNoEEG, and DimEEG). To check whether these values differed between playlists, we performed a one-way repeated-measures analysis of variance (ANOVA) for normally distributed data. Subsequent paired t-tests were used to examine differences between specific playlists. For non-normally distributed data, the Friedman test and the subsequent paired Wilcoxon signed-rank test were used. Normality assumptions were checked using the Shapiro-Wilk test. Similar analyses were performed for all the other items (Table 1). The *p*-values for multiple testing were corrected using Holm’s method (denoted as *p_Holm_*). As measures of effect size, we calculated partial eta-squared for one-way ANOVAs and eta-squared for Friedman tests.

To examine whether the pleasure levels decoded from the EEG differed among playlists, we constructed a Bayesian generalized linear mixed-effects model (GLMM) using the ‘brms’ package in R (ver. 2.16.3)^50-52^. We excluded three additional participants from the analysis because more than 30% of their EEG data were missing during DimNoEEG, resulting in 17 participants. The GLMM was specified using the following formula:

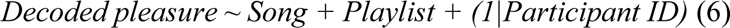

The response variable was the baseline-corrected decoded pleasure from the in-ear EEG (mean per song). The model included fixed effects for Song (the number of songs) and Playlist and a random intercept for each participant to account for individual differences. Playlists were treated as categorical factors, with DimEEG set as the reference level for comparisons. Priors for the model parameters were set to the default non-informative priors provided by brms, except for the intercepts and standard deviations, which were left unspecified. We assumed that the residual errors followed a Student’s t-distribution to accommodate potential outliers and heavier tails in the data. Markov Chain Monte Carlo (MCMC) sampling was performed with the following settings: 4 chains, 2,000 iterations per chain, 1,000 warm-up samples per chain, resulting in 4,000 total post-warm-up samples. The convergence of the MCMC chains was assessed using the potential scale-reduction factor (R-hat). After fitting the GLMM to the data, we conducted pairwise comparisons between playlists using the ‘emmeans’ package (ver. 1.10.3)^53^. We computed the estimated marginal means and performed pairwise contrasts with 95% highest posterior density (HPD) intervals. We evaluated whether the 95% confidence interval of the estimated slope included zero. For subsequent comparisons, we evaluated whether the 95% HPD interval included zero.

Finally, we performed stepwise multiple regression analyses to explore the J-BMRQ and Gold-MSI factors that influence the pleasure of playlists with and without EEG. We conducted the analysis on the same 17 participants as in the previous GLMM. We calculated differences in chills (Chilldiff) and subjective pleasure (Pldiff) between the two EEG playlists (AugEEG minus DimEEG). We then calculated the z-scores of Chilldiff and Pldiff and summed them to obtain the total pleasure score for the EEG playlists (total pleasure EEG). We created a model with 10 factors (five from J-BMRQ and five from Gold-MSI) as explanatory variables, and subjective pleasure or total pleasure EEG as the response variable. A stepwise regression approach was used to determine the optimal model by searching for the lowest Akaike Information Criterion (AIC) value. We also calculated the total pleasure score by subtracting the pleasure score of DimNoEEG from that of AugNoEEG (total pleasure NoEEG: AugNoEEG minus DimNoEEG). Another multiple regression model was constructed with the total pleasure NoEEG score as the response variable, using a similar stepwise method. We used the ‘stepAIC’ function in the ‘MASS’ package (ver. 7.3.54)^54^. To calculate the variance inflation factor (VIF) for the optimized models, we used the ‘vif’ function from the ‘car’ package (ver. 3.1.2)^55^. All tests were two-sided with a significance level of α = 0.05.

### Data availability

The data supporting the findings of this study are available from the Open Science Framework repository at https://osf.io/kx7eq/.

## Acknowledgements

The authors would like to acknowledge Dr. Ernest Mas Herrero for valuable comments on our preliminary results.

## Author contributions

Conceptualization: SK, TE, SF

Data Curation: SK

Formal Analysis: SK

Funding Acquisition: SK, SF

Investigation: SK

Methodology: SK, TE, YS, TI, SF

Project Administration: SF

Software: SK, TI

Supervision: SF

Visualization: SK

Writing – Original Draft Preparation: SK

Writing – Review & Editing: SK, TE, YS, YN, YI, TI, SF

## Competing interests

The authors have read the journal’s policy and have the following competing interests: All authors associated with this research are employed by VIE, Inc. This does not alter our adherence to the journal’s policy of sharing data and materials.

**Materials & Correspondence:** Shinya Fujii

## Extended Data

**Extended Data Fig.1.**
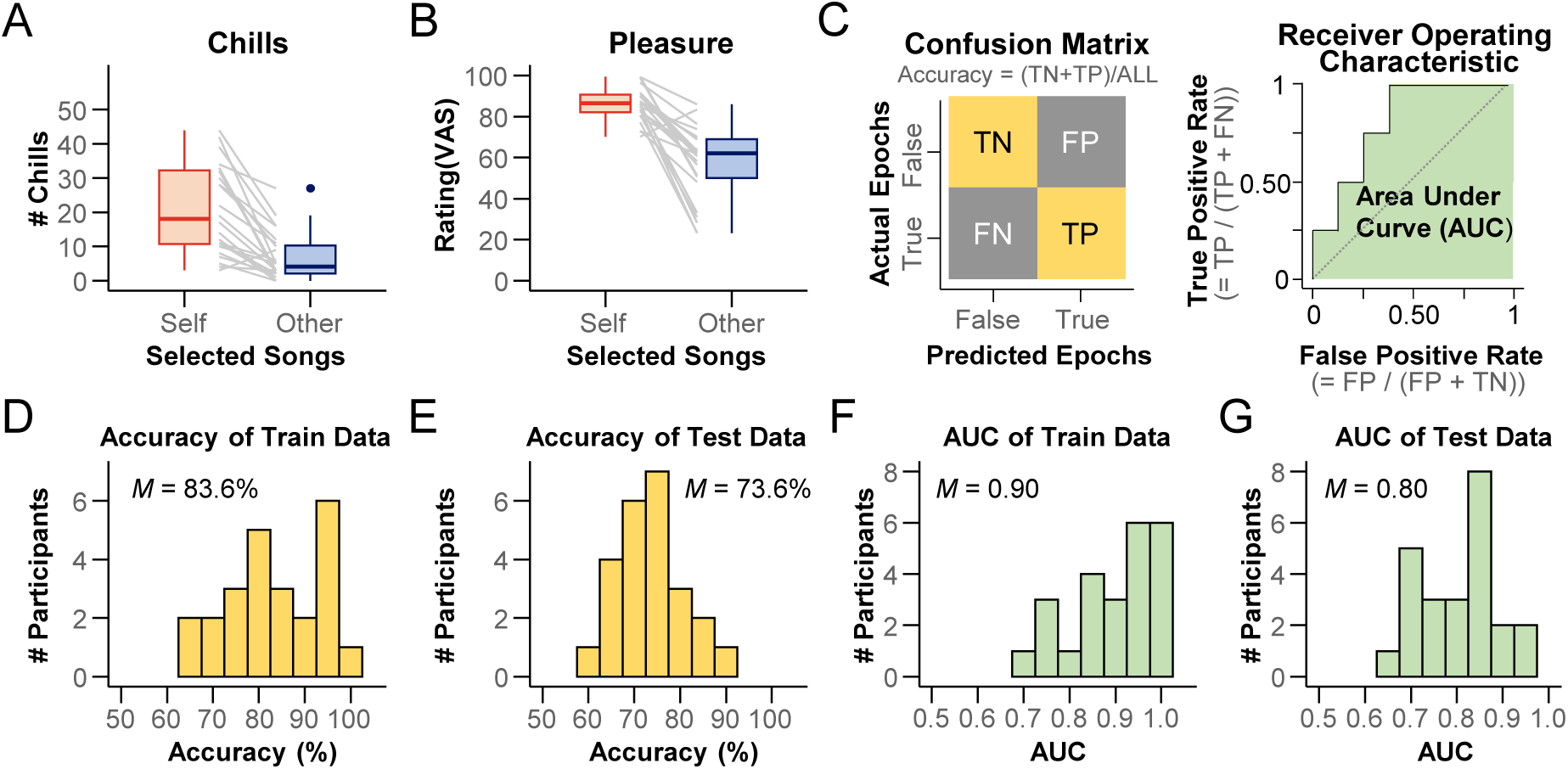
Validation index for a pleasure-predicting model from in-ear EEG (Model 2). A, **B.** Number of chills and pleasure ratings for self-selected and other-selected songs. Except for two participants, participants experienced three or more chills or higher pleasure with self-selected songs than with other-selected songs. **C.** Schematic drawings of confusion matrix (left) and receiver operating characteristic (ROC) curve (right). True and False for the predicted and actual EEG epochs refer to the epochs of self-selected and other-selected songs, respectively. The true Positive (TP), True Negative (TN), False Negative (FN), and False Positive (FP) values were calculated. The area under the curve (AUC) was calculated using the ROC curve. **D, E.** Accuracy of training and test data. The mean accuracies were 83.6% and 73.6%, respectively. **F, G.** AUC of the training and test data. The mean AUCs were 0.90 and 0.80, respectively.

**Extended Data Table 1.** Descriptive statistics (mean ± SD) for the rating values of each item and chills counts, including results of Shapiro-Wilk test.

| Item | AugEEG<br>(*W*-value, *p*-value) | AugNoEEG<br>(*W*-value, *p*-value) | DimNoEEG<br>(*W*-value, *p*-value) | DimEEG<br>(*W*-value, *p*-value) |
| --- | --- | --- | --- | --- |
| Chills | **10.50 ± 9.52**<br>**(0.87, 0.013)** | **9.50 ± 12.15**<br>**(0.75, < 0.001)** | **6.25 ± 7.64**<br>**(0.75, < 0.001)** | **4.90 ± 6.37**<br>**(0.73, < 0.001)** |
| Pleasure | 69.47 ± 20.04<br>(0.92, 0.085) | 60.05 ± 19.60<br>(0.94, 0.285) | 51.06 ± 21.90<br>(0.96, 0.472) | 49.17 ± 20.05<br>(0.913, 0.075) |
| Excitement | 63.27 ± 22.32<br>(0.97, 0.791) | 58.28 ± 21.61<br>(0.96, 0.600) | 52.85 ± 21.00<br>(0.97, 0.718) | 45.24 ± 21.25<br>(0.92, 0.081) |
| Absorption | 69.42 ± 23.34<br>(0.93, 0.141) | 63.47 ± 22.88<br>(0.96, 0.637) | 53.16 ± 22.76<br>(0.98, 0.945) | 50.11 ± 24.38<br>(0.97, 0.753) |
| Arousal | 66.05 ± 20.32<br>(0.97, 0.783) | 58.07 ± 20.58<br>(0.98, 0.899) | 52.79 ± 19.81<br>(0.96, 0.556) | 47.71 ± 27.15<br>(0.96, 0.553) |
| Valence | 68.75 ± 18.30<br>(0.96, 0.504) | **62.34 ± 21.13**<br>**(0.88, 0.016)** | 56.18 ± 17.16<br>(0.98, 0.972) | 56.61 ± 18.73<br>(0.94, 0.282) |
| Preference | 68.24 ± 22.95<br>(0.94, 0.244) | 53.37 ± 23.30<br>(0.99, 0.984) | 43.44 ± 21.67<br>(0.97, 0.665) | 42.79 ± 20.33<br>(0.98, 0.959) |
| Strangeness | 35.36 ± 20.67<br>(0.96, 0.624) | 38.46 ± 20.83<br>(0.97, 0.728) | 42.80 ± 22.95<br>(0.97, 0.832) | 48.52 ± 23.42<br>(0.95, 0.390) |
| Life Purpose | **50.93 ± 24.25**<br>**(0.90, 0.039)** | 48.47 ± 21.84<br>(0.92, 0.080) | **41.33 ± 20.58**<br>**(0.85, 0.005)** | 36.65 ± 19.77<br>(0.93, 0.127) |
| Self-acceptance | 48.21 ± 20.71<br>(0.94, 0.202) | 40.14 ± 22.73<br>(0.91, 0.058) | 43.09 ± 21.13<br>(0.98, 0.859) | 41.18 ± 23.49<br>(0.95, 0.439) |
| Improving relations | 37.63 ± 20.10<br>(0.97, 0.729) | 38.98 ± 21.80<br>(0.93, 0.167) | 28.93 ± 18.49<br>(0.96, 0.522) | 29.93 ± 21.48<br>(0.95, 0.312) |
| Reducing stress | 63.08 ± 19.10<br>(0.99, 0.995) | 53.25 ± 22.65<br>(0.94, 0.215) | 50.86 ± 18.63<br>(0.93, 0.176) | 47.10 ± 21.88<br>(0.95, 0.424) |
| Resilience | 45.38 ± 21.08<br>(0.95, 0.341) | **40.83 ± 20.91**<br>**(0.90, 0.047)** | 38.27 ± 19.60<br>(0.95, 0.38) | 32.72 ± 20.11<br>(0.94, 0.281) |
| Emotion regulation | 45.77 ± 25.26<br>(0.91, 0.069) | 45.39 ± 21.75<br>(0.93, 0.179) | 38.59 ± 16.62<br>(0.93, 0.184) | 32.73 ± 21.81<br>(0.95, 0.327) |
| Seeking help | 40.19 ± 21.21<br>(0.96, 0.542) | **36.90 ± 21.00**<br>**(0.84, 0.004)** | 33.12 ± 22.48<br>(0.93, 0.175) | 31.74 ± 23.45<br>(0.92, 0.096) |

**Note.** Data that did not meet normality assumptions ( $p < 0.05$ ) based on the Shapiro-Wilk test are shown in bold.

**Extended Data Table 2.** Pairwise comparisons for the playlists in rating of Excitement.

| Contrast | Difference<br>[95%CI Lower, Upper] | *t*-value | df | $p_{\text{Holm}}$ |
| --- | --- | --- | --- | --- |
| **AugEEG–DimEEG** | 18.03<br>[5.49, 30.57] | 3.01 | 19 | 0.036 |
| AugEEG–DimNoEEG | 10.42<br>[−0.18, 21.02] | 2.06 | 19 | 0.215 |
| AugEEG–AugNoEEG | 4.99<br>[−9.19, 19.16] | 0.74 | 19 | 0.605 |
| **AugNoEEG–DimEEG** | 13.04<br>[4.37, 21.72] | 3.15 | 19 | 0.032 |
| AugNoEEG–DimNoEEG | 5.43<br>[−5.30, 16.16] | 1.06 | 19 | 0.605 |
| DimNoEEG–DimEEG | 7.61<br>[−3.25, 18.46] | 1.47 | 19 | 0.476 |

**Note.** Contrasts with significant differences from zero ( $p_{\text{Holm}} < 0.05$ ) are highlighted in bold. CI represents confidence interval.

**Extended Data Table 3.**
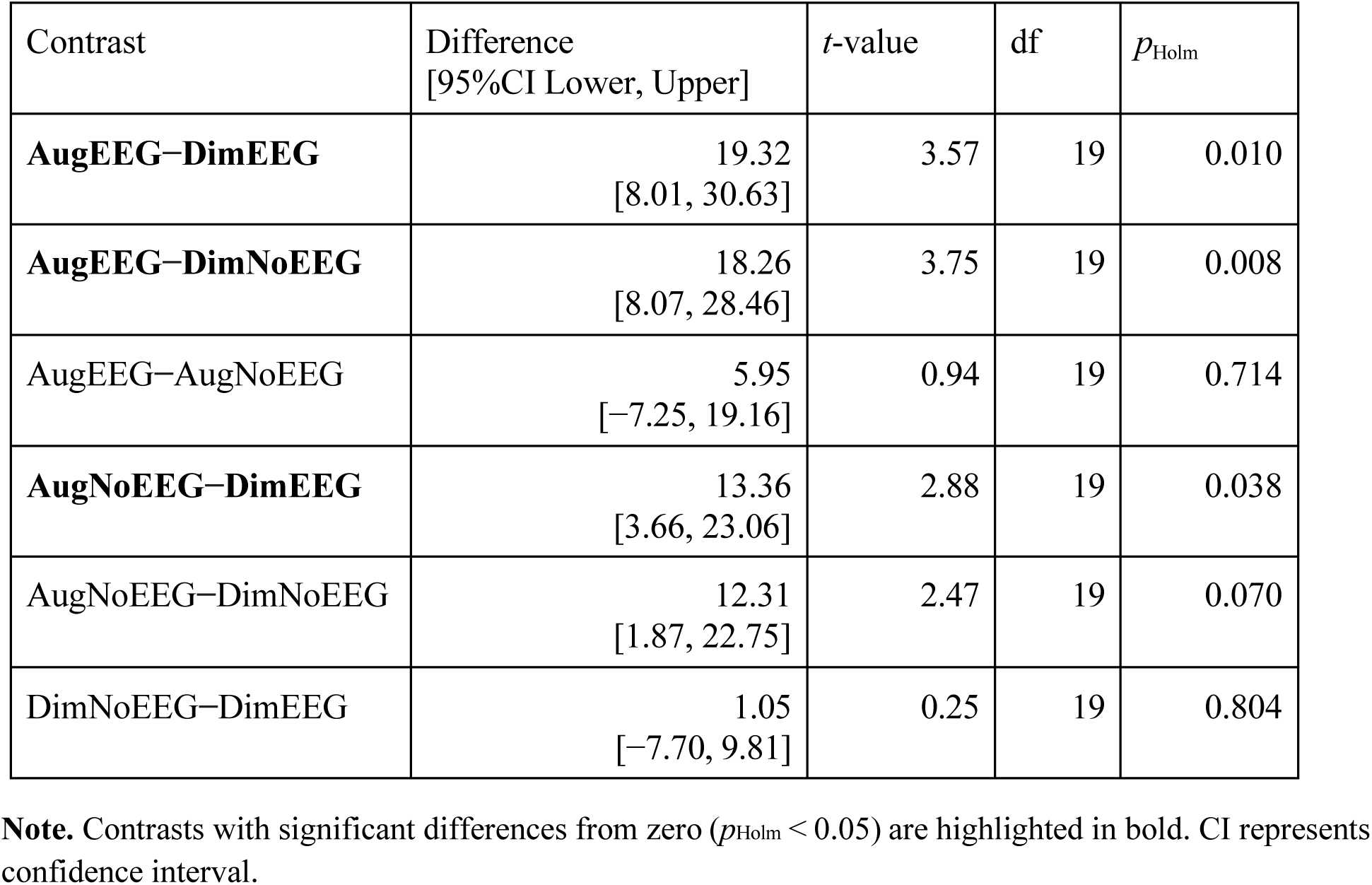
Pairwise comparisons for the playlists in rating of Absorption.

**Extended Data Table 4.** Pairwise comparisons for the playlists in rating of Arousal.

| Contrast | Difference<br>[95%CI Lower, Upper] | *t*-value | df | *p*<sub>Holm</sub> |
| --- | --- | --- | --- | --- |
| **AugEEG–DimEEG** | 18.34<br>[5.70, 30.98] | 3.04 | 19 | 0.041 |
| AugEEG–DimNoEEG | 13.26<br>[3.21, 23.32] | 2.76 | 19 | 0.062 |
| AugEEG–AugNoEEG | 7.98<br>[0.42, 15.55] | 2.21 | 19 | 0.159 |
| AugNoEEG–DimEEG | 10.36<br>[–1.58, 22.30] | 1.82 | 19 | 0.256 |
| AugNoEEG–DimNoEEG | 5.28<br>[–5.63, 16.19] | 1.01 | 19 | 0.496 |
| DimNoEEG–DimEEG | 5.07<br>[–3.83, 13.98] | 1.19 | 19 | 0.496 |

**Note.** Contrasts with significant differences from zero ( $p_{\text{Holm}} < 0.05$ ) are highlighted in bold. CI represents confidence interval.

**Extended Data Table 5.**
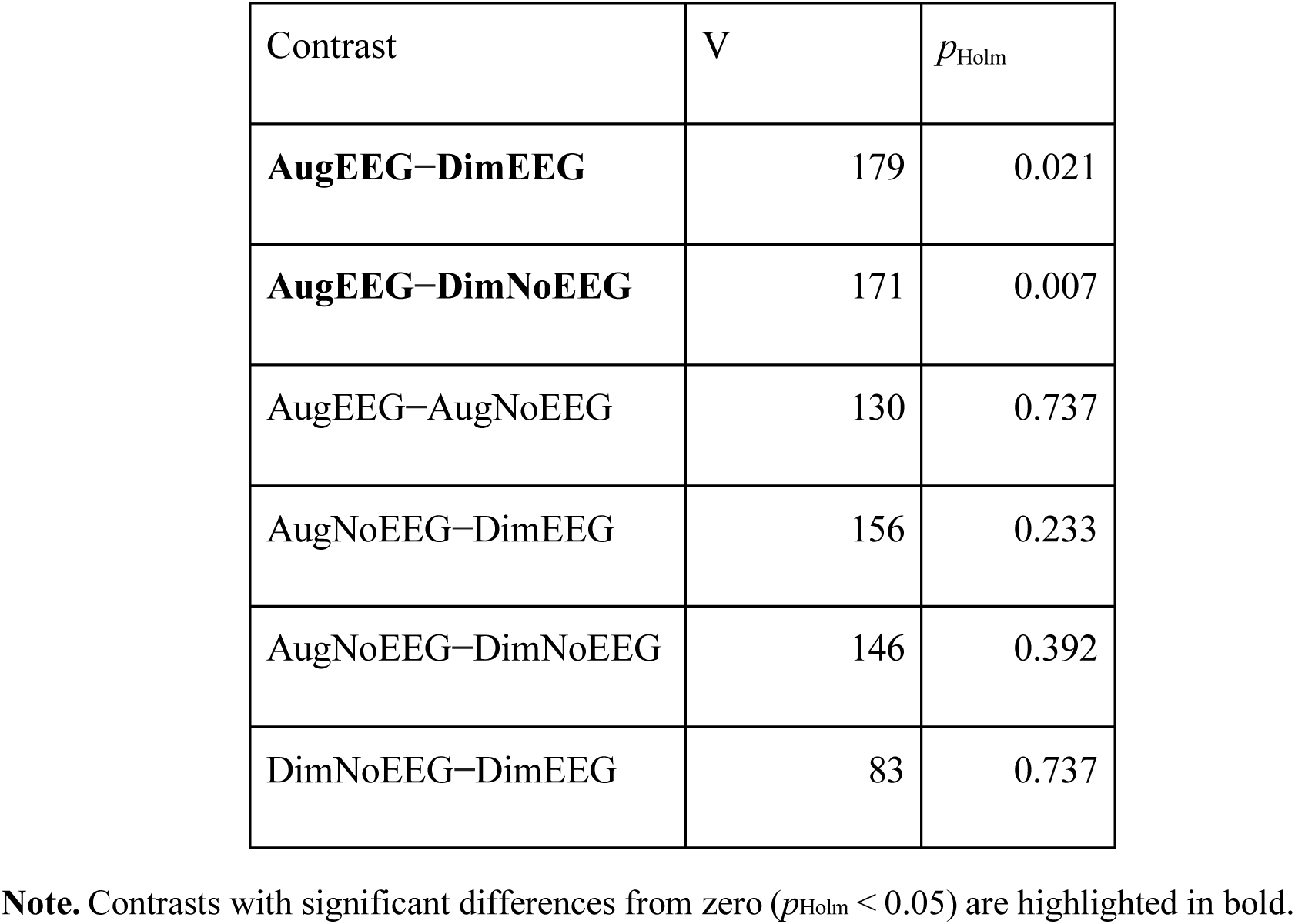
Pairwise comparisons for the playlists in rating of Valence.

**Extended Data Table 6.** Pairwise comparisons for the playlists in rating of Preference.

| Contrast | Difference<br>[95%CI Lower, Upper] | *t*-value | df | $p_{\text{Holm}}$ |
| --- | --- | --- | --- | --- |
| **AugEEG–DimEEG** | 25.45<br>[15.58, 35.32] | 5.40 | 19 | < 0.001 |
| **AugEEG–DimNoEEG** | 24.80<br>[14.33, 35.28] | 4.96 | 19 | < 0.001 |
| AugEEG–AugNoEEG | 14.87<br>[1.65, 28.09] | 2.35 | 19 | 0.118 |
| AugNoEEG–DimEEG | 10.58<br>[–1.38, 22.54] | 1.85 | 19 | 0.239 |
| AugNoEEG–DimNoEEG | 9.93<br>[–2.50, 22.36] | 1.67 | 19 | 0.239 |
| DimNoEEG–DimEEG | 0.65<br>[–8.28, 9.58] | 0.15 | 19 | 0.881 |

**Note.** Contrasts with significant differences from zero ( $p_{\text{Holm}} < 0.05$ ) are highlighted in bold. CI represents confidence interval.

**Extended Data Table 7.** Pairwise comparisons for the playlists in rating of Life purpose.

| Contrast | V | $p_{\text{Holm}}$ |
| --- | --- | --- |
| **AugEEG–DimEEG** | 147 | 0.034 |
| AugEEG–DimNoEEG | 142 | 0.060 |
| AugEEG–AugNoEEG | 109 | 0.790 |
| AugNoEEG–DimEEG | 127 | 0.295 |
| AugNoEEG–DimNoEEG | 101 | 0.790 |
| DimNoEEG–DimEEG | 88 | 0.790 |

**Note.** Contrasts with significant differences from zero ( $p_{\text{Holm}} < 0.05$ ) are highlighted in bold.

**Extended Data Table 8.** Pairwise comparisons for the playlists in rating of Reducing stress.

Pairwise comparisons for the playlists in rating of Reducing stress
| Contrast | Difference<br>[95%CI Lower, Upper] | *t*-value | df | *p*<sub>Holm</sub> |
| --- | --- | --- | --- | --- |
| **AugEEG–DimEEG** | 15.98<br>[6.42, 25.53] | 3.50 | 19 | 0.012 |
| **AugEEG–DimNoEEG** | 12.22<br>[5.19, 19.25] | 3.64 | 19 | 0.011 |
| AugEEG–AugNoEEG | 9.83<br>[–2.62, 22.28] | 1.65 | 19 | 0.460 |
| AugNoEEG–DimEEG | 6.15<br>[–6.88, 19.17] | 0.99 | 19 | 1.000 |
| AugNoEEG–DimNoEEG | 2.39<br>[–9.39, 14.17] | 0.43 | 19 | 1.000 |
| DimNoEEG–DimEEG | 3.75<br>[–7.43, 14.94] | 0.70 | 19 | 1.000 |

**Note.** Contrasts with significant differences from zero ( $p_{\text{Holm}} < 0.05$ ) are highlighted in bold. CI represents confidence interval.

**Extended Data Table 9.** Estimation of a Bayesian GLMM fitted to decoded pleasure.

Estimation of a Bayesian GLMM fitted to decoded pleasure
| Variable | Estimated Slope<br>[95%CI Lower, Upper] | SE | R-hat | bulk-ESS | tail-ESS |
| --- | --- | --- | --- | --- | --- |
| (Intercept) | − 0.034<br>[−0.077, 0.009] | 0.022 | 1.006 | 637 | 1255 |
| **Song** | −0.009<br>[−0.013, −0.005] | 0.002 | 1.000 | 8206 | 3190 |
| **Condition**<br>**(AugEEG – DimEEG)** | 0.034<br>[0.011, 0.057] | 0.012 | 1.002 | 2190 | 2703 |
| Condition<br>(AugNoEEG – DimEEG) | −0.003<br>[−0.026, 0.020] | 0.012 | 1.001 | 2303 | 2614 |
| Condition<br>(DimNoEEG – DimEEG) | 0.020<br>[−0.003, 0.044] | 0.012 | 1.001 | 2327 | 2447 |

**Note.** Variables for which the 95% confidence interval (CI) of the estimated slope did not include zero are highlighted in bold. SE stands for standard error. R-hat indicates the potential scale reduction factor, which assesses the convergence of MCMC chains. Bulk-ESS represents the bulk effective sample size, which indicates the number of effectively independent samples in the central part of the posterior distribution. Tail-ESS represents the tail effective sample size, which indicates the number of effectively independent samples in the tails of the posterior distribution.

**Extended Data Table 10.**
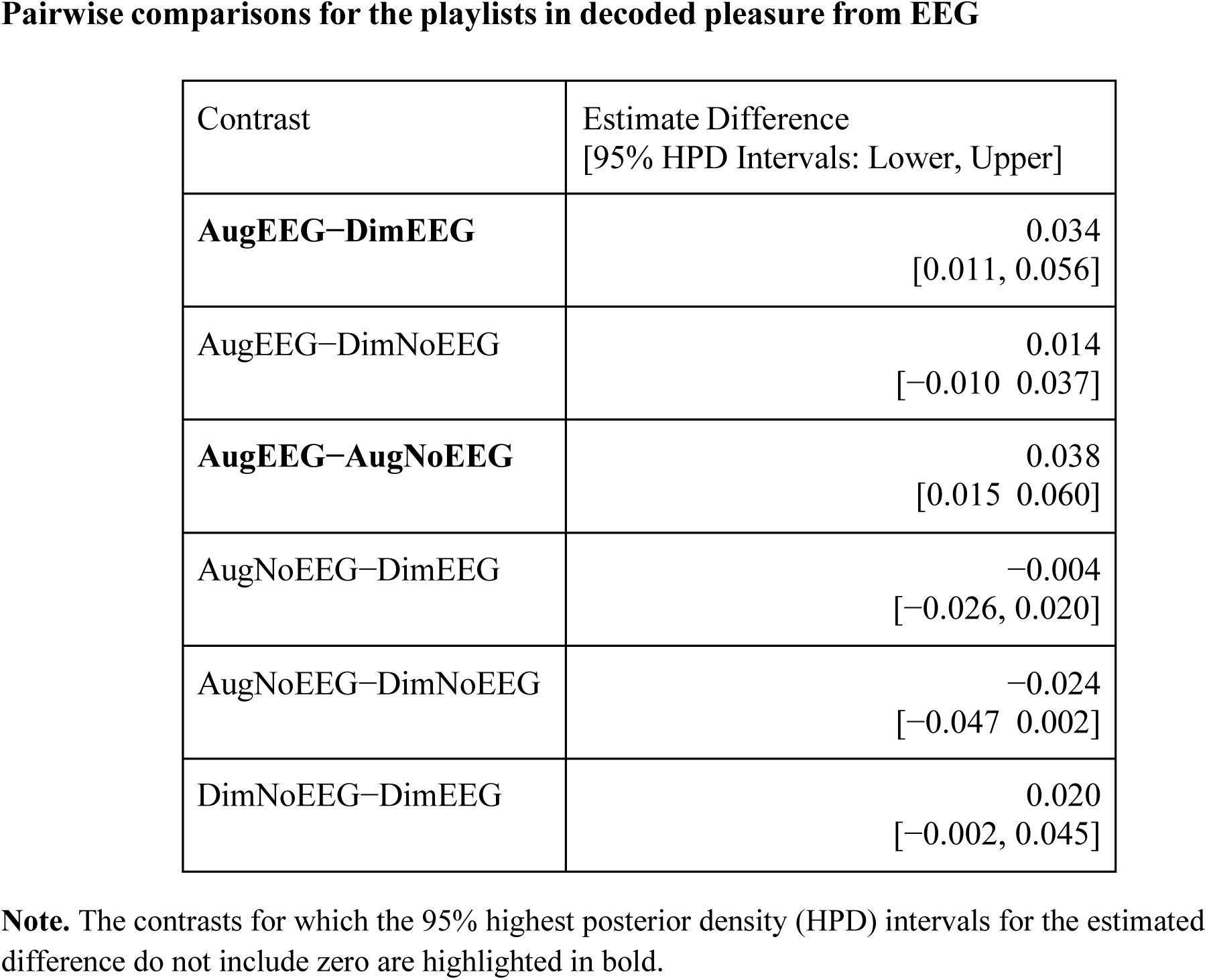
Pairwise comparisons for the playlists in decoded pleasure from EEG.

Pairwise comparisons for the playlists in decoded pleasure from EEG
| Contrast | Estimate Difference<br>[95% HPD Intervals: Lower, Upper] |
| --- | --- |
| **AugEEG–DimEEG** | 0.034<br>[0.011, 0.056] |
| AugEEG–DimNoEEG | 0.014<br>[−0.010 0.037] |
| **AugEEG–AugNoEEG** | 0.038<br>[0.015 0.060] |
| AugNoEEG–DimEEG | −0.004<br>[−0.026, 0.020] |
| AugNoEEG–DimNoEEG | −0.024<br>[−0.047 0.002] |
| DimNoEEG–DimEEG | 0.020<br>[−0.002, 0.045] |

**Note.** The contrasts for which the 95% highest posterior density (HPD) intervals for the estimated difference do not include zero are highlighted in bold.

**Extended Data Table 11.**
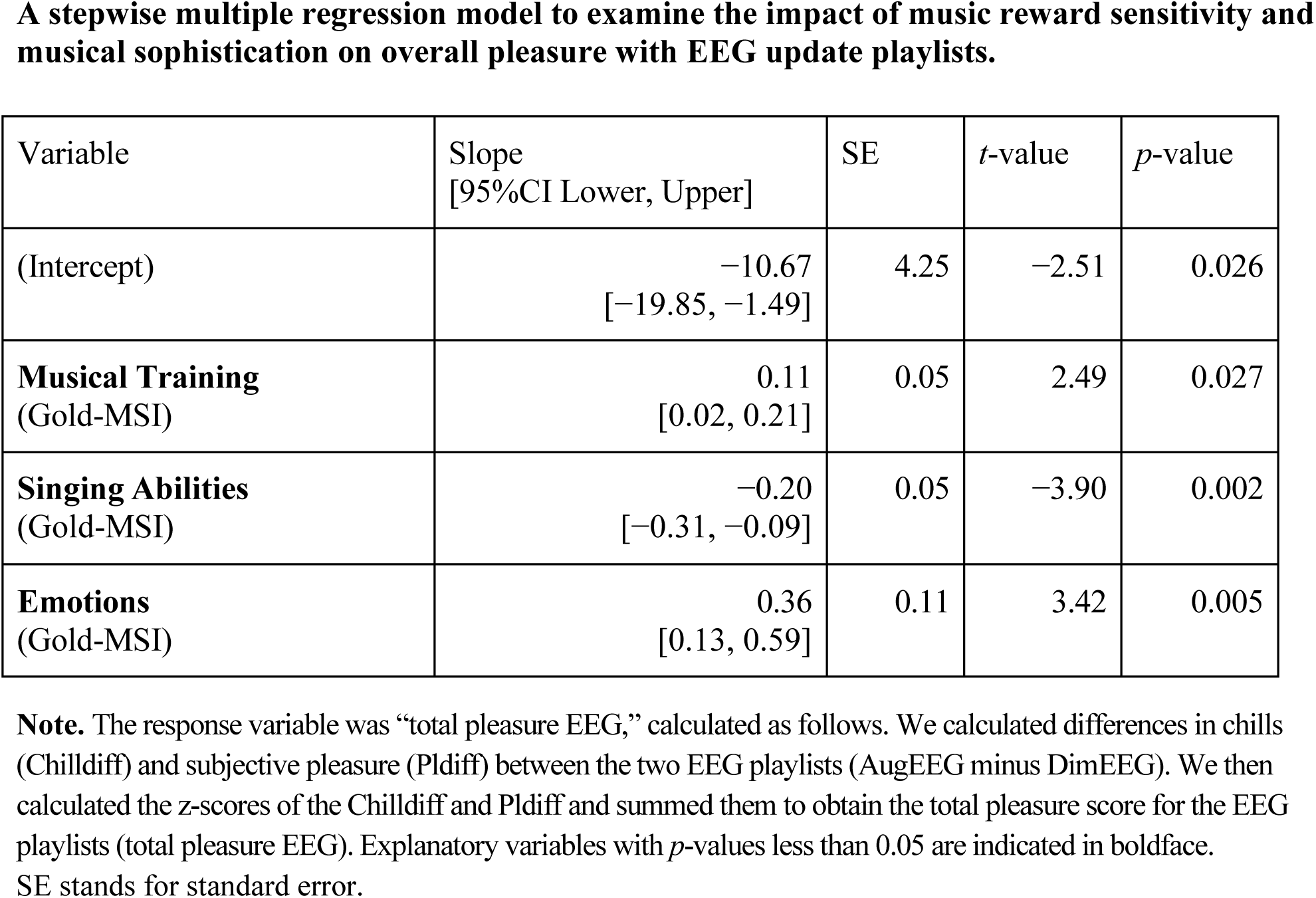
A stepwise multiple regression model to examine the impact of music reward sensitivity and musical sophistication on overall pleasure with EEG update playlists.

| Variable | Slope<br>[95%CI Lower, Upper] | SE | *t*-value | *p*-value |
| --- | --- | --- | --- | --- |
| (Intercept) | −10.67<br>[−19.85, −1.49] | 4.25 | −2.51 | 0.026 |
| **Musical Training**<br>(Gold-MSI) | 0.11<br>[0.02, 0.21] | 0.05 | 2.49 | 0.027 |
| **Singing Abilities**<br>(Gold-MSI) | −0.20<br>[−0.31, −0.09] | 0.05 | −3.90 | 0.002 |
| **Emotions**<br>(Gold-MSI) | 0.36<br>[0.13, 0.59] | 0.11 | 3.42 | 0.005 |

**Note.** The response variable was “total pleasure EEG,” calculated as follows. We calculated differences in chills (Chilldiff) and subjective pleasure (Pldiff) between the two EEG playlists (AugEEG minus DimEEG). We then calculated the z-scores of the Chilldiff and Pldiff and summed them to obtain the total pleasure score for the EEG playlists (total pleasure EEG). Explanatory variables with *p*-values less than 0.05 are indicated in boldface. SE stands for standard error.

**Extended Data Table 12.** A stepwise multiple regression model to examine the impact of music reward sensitivity and musical sophistication on overall pleasure from playlists without EEG updates.

| Variable | Slope<br>[95%CI Lower, Upper] | SE | *t*-value | *p*-value |
| --- | --- | --- | --- | --- |
| (Intercept) | 6.14<br>[−1.15, 13.43] | 3.37 | 1.82 | 0.092 |
| **Mood Regulations**<br>**(BMRQ)** | −0.25<br>[−0.39, −0.10] | 0.07 | −3.74 | 0.002 |
| Sensory Motors<br>(BMRQ) | 0.05<br>[−0.04, 0.15] | 0.04 | 1.29 | 0.219 |
| Perceptual Abilities<br>(Gold-MSI) | 0.09<br>[−0.05, 0.23] | 0.07 | 1.33 | 0.207 |

**Note.** The response variable was “total pleasure NoEEG,” calculated as follows. We calculated the differences in chills (Chilldiff) and subjective pleasure (Pldiff) between the two playlists without using the EEG (AugNoEEG minus DimNoEEG). We then calculated the z-scores of Chilldiff and Pldiff and summed them to obtain the total pleasure score (total pleasure NoEEG). Explanatory variables with *p*-values less than 0.05 are indicated in boldface. SE stands for standard error.

**Extended Data Table 13.** Information on self-selected songs and their artists or composers.

Information on self-selected songs and their artists or composers.
| Song 1 | Song 2 | Song 3 |
| --- | --- | --- |
| Melody / Koji Tamaki | Kaze / Kobukuro | Towa ni tomoni / Kobukuro |
| Neon / John Mayer | Bad / Christopher | Finesse / Bruno Mars |
| Symphony No.1 / J. Brahms | Violin Sonata / R. Strauss | Symphony No.6<br>/ P. Tchaikovsky |
| All My Life / Lil Durk | Imagine / Snoop Dogg | B2B / BAD HOP |
| Love Theme<br>/ Ennio Morricone | Hanataba wo Kimi ni<br>/Hikaru Utada | Intermezzo Sinfonico<br>/P. Mascagni |
| So Sick / Ne-Yo | Ur Perfect I Hate It (Lenno<br>Remix) / Mickey Valen | Green Green Grass (Sam Feldt<br>remix) / George Ezra |
| astrid / glaive | Gold / Hiakru Utada | Vivienne Westwood / lileffort |
| La traviata Act I / G. Verdi | This Is Me / Keala Settle | Locus iste / A. Bruckner |
| Flowers (feat. Nori)<br>/in love with a ghost | Repeatedly - Video Edit<br>/Ametsub | Florence / Kevin Penkin |
| One Last Kiss / Hikaru Utada | Kiss & Cry / Hikaru Utada | Traveling / Hikaru Utada |
| Planetarium / Justin Hurwitz | Hannah Song / Sam Wilkes | Reve Rouge / Cirque du Solei |
| Piano Concerto in G major<br>/ M. Ravel | McDonald Romance<br>/ King Gnu | Out of This World<br>/ Masato Kumoi |
| Whenever I GO<br>/ Jacob Collier | Witness Me / Jacob Collier | Burn The Witch / Radiohead |
| Geek U.S.A.<br>/ Smashing Pumpkins | Readymade<br>/ Red Hot Chili Peppers | My Hero / Foo Fighters |
| rose / KID FRESINO | Linkin Park / Numb | Dancing Boy / OKAMOTO'S |
| Reimei, / Δ | First Rain / Kontinuum | Saikai / Seigen Tokuzawa |
| Utsukushiu hi<br>/SUPER BEAVER | Symphony No.5<br>/ P. Tchaikovsky | Seikai / RADWIMPS |
| Zakir / John McLaughlin | Black is the color of my true<br>love's hair / Nina Simone | Ex-Factor / Lianne La Havas |
| youth ft. Hakushi Hasegawa<br>/ KID FRESINO | Yumeutsutsu / Kenshi Yonezu | Pale Blue / Kenshi Yonezu |
| Ode to Joy / L. Beethoven | Sonate für Klavier Nr.17<br>D-Dur D 850 Op.53<br>/ F. Schubert | FAMILIA<br>/ MILLENNIUM PARADE |
| Snow White<br>/ Keisuke Kuwata | So Close / Jon Mclaughlin | Part of Your World<br>/ Halle Bailey |
| The others / MIYAVI | Uminari / Kodo | Zoku / Kodo |
| Synthesis / tn-shi | Golden Hour / Soflynaily | STYX HELIX<br>/ MYTH & ROID |
| We Are / ONE OK ROCK | Tremolo / RADWIMPS | Make A Wish /<br>ELLEGARDEN |

**Note.** Each row shows self-selected songs provided by each participant. There were no duplicates of the songs among the participants.

